# Drought Response and Recovery in Wild, Feral, and Domesticated *Brassica rapa*: Implications for Crop Climate Resilience

**DOI:** 10.64898/2026.11.06.764879

**Authors:** Aleca M. Borsuk, Gabriella I. Quartuccia, Chase Talpins, Grace Angert, Rachel Warner, Luis R. Vega, Meredith Leahy, Rhea Mittal, Leeann C. Daybydeen, Isabella Grippo, Julie Bichler, Elisabeth Moore, Reem Salha, Kelly Ivanovski, Anna Ellingboe, Giulia Mattalia, Makenzie Mabry, Mauricio Diazgranados, Steven J. Franks, Alex C. McAlvay

**Author notes:** These authors contributed equally to the work (Aleca M. Borsuk, Gabriella I. Quartuccia). **Corresponding Author**: Aleca Borsuk.

## Abstract

Drought poses a serious threat to agricultural production and food security. Wild relatives of crops, including pre-domesticated and feral varieties, contain genetic diversity that can serve as a breeding resource for increasing desirable crop traits such as drought resilience, but little is known about the relative ability of wild varieties to cope with climate change in most species. In this study, we investigated the drought response of wild, feral, and domesticated varieties of *Brassica rapa*, an important crop and model species. Twenty-five accessions were subjected to experimental drought and recovery, and variation in traits such as biomass retention, photosynthetic rate, and water use efficiency were quantified. We found several wild accessions of *B. rapa* had high biomass retention under drought and full recovery of maximum assimilation rate after rewatering, challenging the assumption that feralization produces stress-resilient phenotypes and suggesting that populations of wild crop progenitors are especially promising reservoirs of drought-response diversity. Our results also showed taproot biomass was positively associated with water use efficiency. Our findings provide candidate traits and genotypes for agricultural breeding aimed at increasing drought resilience in domesticated *B. rapa* cultivars and give insights into evolution under domestication and feralization in the context of climate change.

**Highlight:** Drought-resilient traits and genotypes were identified in wild, feral, and domesticated accessions of *Brassica rapa*, providing insight into evolution under domestication and feralization in the context of climate change.

## Introduction

Drought is one of the most severe constraints on crop productivity worldwide, and its impacts are becoming increasingly problematic due to climate change (Trenberth et al., 2014; Kogan et al., 2019; Begna and Wakweya, 2025). Despite its importance to agriculture, drought resilience, i.e. a plant’s ability to recover after experiencing drought stress, remains a challenging trait to understand and breed for due to the fact that it involves multiple physiological, morphological, and developmental processes (Khadka et al., 2020). These responses are context-dependent and often controlled by multiple genes, making it difficult to select for drought-resilient genotypes using conventional breeding approaches (Mir et al., 2012).

Crop wild relatives (CWRs) are potentially valuable resources for breeding crops with enhanced drought resilience due to relatively high genetic diversity and traits that enable survival under harsh environmental conditions (Zhang et al., 2017; Jovovic et al., 2020). CWRs offer reservoirs of genetic diversity that can contribute to crop resilience under climate change (Dempewolf et al., 2014; Brozynska et al., 2016). In recent years, interest has grown in leveraging these populations not only as a source of traits but also through novel approaches such as *de novo* domestication—where wild organisms are domesticated from scratch (Fernie and Yan, 2019). Despite this potential, there is limited information on how domesticated crops and their wild forms vary in terms of physiological traits such as photosynthetic rate, water use efficiency, and stomatal conductance (Lei et al., 2024).

Feral crops, which are formerly domesticated populations of plants that have escaped cultivation and form free-living populations, are also potentially underutilized pools of adaptive diversity for breeding stress resilience, as these populations often persist in disturbed or marginal habitats, suggesting they may possess valuable stress resilience traits (Ellstrand et al., 2010; Mabry et al., 2023). *De novo* re-domestication, which uses feral populations as raw material to breed new crops, has also been proposed as another way to leverage adaptive diversity (Pisias et al., 2022). Yet, feralization is relatively understudied: while studies have demonstrated the presence of “domestication syndromes,” which are morphological, physiological, and developmental traits that commonly arise in groups of plants (and animals) under domestication (Meyer et al., 2012; Rendón-Anaya and Herrera-Estrella, 2018), analogous “feralization syndromes” are comparatively less well characterized, especially outside of work on the primary model for feral plant biology, weedy rice (Gressel, 2005; Mabry et al., 2021; Mabry et al., 2023). Pathways to feralization do not necessarily follow the reverse of domestication, making it important to understand feral plant physiology and genetics as unique phenomena (Niego and Benítez-Burraco, 2022).

The physiological mechanisms underlying drought responses remain poorly understood across domestication status, i.e., from wild to domesticated to feral, in most crop systems (Milla et al., 2015). In particular, there is a lack of direct experimental studies that compare plant responses across domestication status under controlled drought conditions, limiting our understanding of how domestication and ferality shape stress adaptation and our ability to predict the presence of beneficial traits in wild and feral crop relatives.

*Brassica rapa* presents a powerful model system for understanding drought response from the perspective of evolutionary ecology across domestication status. *B. rapa* includes a range of crop types selected for different target organs, including leafy vegetables (e.g., bok choy), root crops (e.g., turnip), and oilseeds (e.g., yellow sarson). The genus *Brassica* includes a number of crops of major economic and nutritional importance, with many of them forming what is called the “Triangle of U” with the diploid species *Brassica rapa*, *Brassica oleracea*, and *Brassica nigra* on the vertices, and allotetraploids of each combination of the three diploids forming the sides (Nagaharu and Nagaharu, 1935). However, *B. rapa* in particular has emerged as a valuable model for multiple areas of plant biology due to its phylogenetic proximity to the plant model *Arabidopsis thaliana*, genomic resources, rapid cycling forms, and role as a genome donor to the well-studied allopolyploid relatives *Brassica napus* and *Brassica juncea* (McAlvay et al., 2021). Drought poses a major constraint for many *Brassica* crops, with significant yield losses reported under water-limited conditions (Chikkaputtaiah et al., 2017). Field mustard, *B. rapa*, has also emerged as an important model for understanding rapid evolutionary responses to drought (Franks, 2011; Franks et al., 2016; Hamann et al., 2018), including early flowering in wild populations as a mechanism for drought escape (Franks, 2011).

To date, comparative studies in *B. rapa* across the entire wild-feral-domestication spectrum have been hampered by a lack of clarity over which populations are feral vs. truly wild (McAlvay et al., 2017). In a recent advance, population genetic work has clarified the domestication status in 653 accessions of *B. rapa*, identifying wild lineages from the Caucasus, Siberia, Italy, and Algeria. The study used genotyping-by-sequencing and habitat suitability mapping to understand genetic relationships, diversity, admixture, and structure to distinguish their evolutionary status. This set of resolved wild, feral, and domesticated populations provides an unusually clear system to compare drought response across domestication status (McAlvay et al., 2021).

In this study, we took advantage of these newly resolved genetic relationships to understand the impact of domestication status on *B. rapa* response to drought. Specifically, we aimed to understand if wild, domesticated, and feral *B. rapa* differ in drought response, and if so, what physiological traits support recovery from drought, i.e., resilience. To inform future crop breeding efforts, we also aimed to understand if specific accessions of *B. rapa* had elevated drought resilience. Finally, we sought to understand if there was variation in drought response across crop types (e.g., leafy vs root vs oilseed), including the impact of fleshy taproots on plant water use efficiency. To address these questions, we carried out a greenhouse drought and recovery experiment comparing 25 accessions of *B. rapa* across domestication status and crop types, incorporating gas exchange, chlorophyll fluorescence, and biomass data. The overall goal of this work was to contribute to our fundamental understanding of adaptive drought response in wild, domesticated, and feral plants, and to inform agricultural breeding efforts aimed at increasing drought resilience in *B. rapa* crops.

## Materials and Methods

### Plant material and growth conditions

A set of 25 accessions of *B. rapa* was selected to represent a broad range of geographies, domestication statuses, and morphotypes. This included nine wild accessions, eight feral accessions, and eight crop accessions (three turnips, two oilseeds, and three ‘leafy’, i.e. bok choy/choy sum). All accessions were selected as a subset of total accessions sequenced for a previous study (McAlvay et al., 2021) to identify their status as wild, feral, or the specific crop. Fresh seeds were obtained from the Leibniz Institute of Plant Genetics and Crop Plant Research, United States Department of Agriculture, and UK Vegetable Genebanks. Plants were grown in 6-inch standard pots in a commercial high-porosity soil (PRO-MIX HP Mycorrhizae; Premier Tech Horticulture, Québec, Canada) containing 65-75% sphagnum moss, perlite, limestone, a wetting agent, a starter nutrient, and a mycorrhizal component (*Rhizophagus irregularis*), amended with compost and RootShield granular biofungicide (*Trichoderma harzianum* strain T-22) at the Nolen Greenhouses at the New York Botanical Garden.

Potted plants were arranged in a randomized block design during the experimental phase (Fig. S1). Each block consisted of one individual of each of the 25 accessions. A total of ten blocks (A through J) were established, five of which were subjected to experimental drought and recovery (i.e., ‘drought’ treatment) and five of which were consistently well-watered (i.e., ‘control’ treatment).

All plants were watered by hand to maintain soil hydration throughout germination and establishment. Starting at the onset of the experimental phase, blocks A-E were designated as the control treatment, and watered once per week by saturating the soil completely via timed (10-min) drip irrigation. Blocks F-J were designated as the drought treatment and only watered in week 0 (start of the experiment) and then again in week 3 (recovery watering), to full saturation of the soil via timed (10-min) drip irrigation (Table S2).

### Gas exchange and chlorophyll fluorescence

Leaf gas exchange was measured 1-3 days immediately following plant watering (due to the number of replicates, gas exchange measurements could not be completed on all plants in a single day). Rate of maximum CO_2_ assimilation (A_max_) was measured using a LI-6800 portable gas exchange system (Licor Inc, Lincoln, NE, USA); leaves were placed in the chamber during the hours of 9 am to 2 pm and allowed to acclimate to the following conditions until they reached a steady state: PPFD = 800 µmol m^-2^ s^-1^, CO_2_ = 400 ppm, RH = 37%, VPD = 1.6 kPa, T_leaf_ = 25 °C. The light intensity for all A_max_ measurements was standardized at 800 PPFD, to reduce photoinhibition during drought and accommodate a variety of accessions adapted to different light environments. Stomatal conductance to water (g_sw_) was monitored simultaneously with assimilation rate to calculate instantaneous water use efficiency (WUE = A_max_/g_sw_). To estimate photosynthetic efficiency as an indicator of stress, the quantum yield of photosystem II (YII) was measured pre-dawn (between ∼3 am to ∼6 am) the day after watering on dark-adapted plants using a MINI-PAM-II fluorometer (WALZ GmbH, Effeltrich, Germany). For all gas exchange and chlorophyll fluorescence data, measurements were obtained from 1 leaf per individual and replicated on 5 individuals per accession for each treatment (drought and control). Measurements were obtained from leaves at the same developmental stage (leaf 4-5 in weeks 0-1, and leaf 5-6 in weeks 2-3).

### Soil moisture

Soil moisture was tested synchronously with gas exchange measurements using a Hydrosense II soil moisture sensor (Campbell Scientific, Logan, Utah) to ensure the drought treatment was effective and comparable across domestication status and crop type (Fig. S2). The moisture probe was inserted vertically at the midpoint between the stem and pot wall and held in place until the measurement stabilized. For each sampling event, two measurements were taken at opposite sides of the pot. The two volumetric water content readings were averaged to determine the mean soil moisture.

### Biomass

Plant biomass was collected for all individuals at the conclusion of the growth experiment and partitioned into vegetative, hypocotyl (i.e., fleshy taproot), and root tissues. Roots were separated from soil and other particulates by first submerging the root ball in water to remove bulk soil content, then subsequent washing of the roots under running tap water. All tissue samples were dried in paper bags at 60°C in a gravity oven (Fisherbrand™, Waltham, Massachusetts, USA). Dry weight of each sample was weighed using a 0.001 g balance (Sartorius, Göttingen, Germany).

### Statistical Analysis

#### Biomass retention

Total plant biomass (the sum of vegetative, hypocotyl, and root tissue mass) was calculated for each accession in drought vs. control treatments. Biomass retention was calculated for each accession as the ratio of total plant biomass in the drought treatment to total plant biomass in the control treatment (Biomass retention = Drought biomass / Control biomass). Values >1 indicate relative biomass gain in the drought treatment compared to the control treatment, while values <1 indicate biomass loss in the drought treatment compared to the control treatment.

#### Comparison of biomass retention group means

Linear models were used to test the effect of domestication status (wild, n =9; feral, n = 8; domesticated, n = 8) on biomass retention. Model assumptions were checked with Shapiro-Wilk and Levene’s test. Estimated marginal means were calculated via the *emmeans* package in R (Lenth and Piaskowski, 2025) and pairwise comparisons were run between categories using Šidák adjustment for multiple comparisons.

#### Physiological response models

Linear mixed effects models (LMEM) were run via the “lmer” command from the *lme4* package in R (Bates et al., 2015) to test how A_max_, YII, water use efficiency, or stomatal conductance (physiological response; Tables S3-S5) varied with fixed and random effects. LMEM included treatment, week, and domestication type as fixed effects with all two- and three-way interactions between terms, as well as the random effects of block and plant ID nested within accession. The *emmeans* package was used to make pairwise comparisons between groups via Tukey adjustment. LMEM were run to compare cultivar physiological response at the weekly level, as well as across two weeks at a time to compare the rate of change in physiological response between the two treatments. LMEM report Tukey-adjusted p-values. Linear models were additionally run using “lm” regression from the *stats* package in R (R Core Team, 2025) to investigate the relationship between water use efficiency and mean taproot biomass.

## Results

### Biomass Retention

The drought treatment significantly reduced biomass overall, as expected, with plants in the drought treatment having on average 78.6% of the biomass of plants in the watered treatment (t_(48)_ = -2.64, p = 0.011). However, there was substantial variation among varieties, with some having greater biomass under drought than control conditions. Specifically, a biomass retention index (drought biomass / control biomass) greater than 1, which indicates greater biomass in drought than control conditions (overcompensation), occurred in three wild accessions native to the country Georgia, a domesticated accession from Finland, and a feral accession from Guatemala (Fig. 1A; Table S1). In contrast with these highly drought resilient varieties, some were very drought sensitive, including two domesticated and one feral variety that had less than half of the biomass under drought compared with control conditions (Fig. 1A). Comparing accessions by domestication status, biomass retention was significantly higher in wild accessions (mean = 1.05 ± 0.089) than in domesticated accessions (mean = 0.723 ± 0.089; t_(22)_ = -2.60, p = 0.049), with feral plants (mean = 0.799 ± 0.101) not differing from either wild (t_(22)_ = -1.86, p = 0.212) or domesticated accessions (t_(22)_ = -0.57, p = 0.924; Fig. 1B). When domesticated accessions were further compared by crop type (e.g., leafy vs root vs oilseed), and after adjusting for multiple comparisons, no crop type differed significantly in biomass retention (Fig. S3).

**Figure 1.**
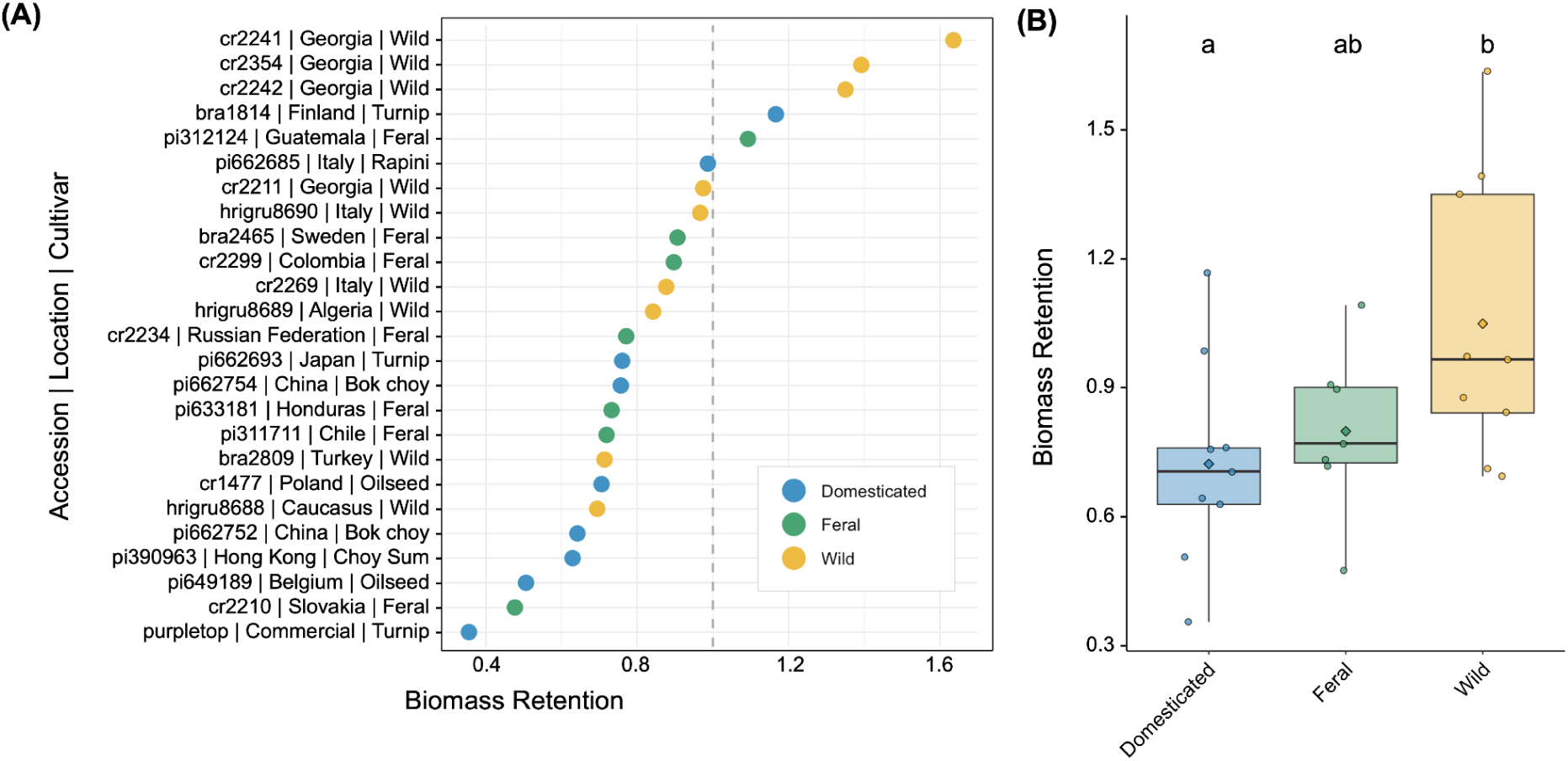
Biomass retention (drought biomass/control biomass) comparisons across 25 accessions of domesticated, feral, and wild *Brassica rapa*. (A) Ranking of accessions by biomass retention; dashed line indicates a biomass retention value of one. (B) Comparison of biomass retention by domestication status. Box upper and lower boundaries represent the 75^th^ and 25^th^ percentiles, respectively, and lines across boxes represent the group median. Whiskers extend from the upper and lower quartiles to the group maximum and minimum, respectively. Group means not sharing a letter are significantly different by pairwise comparisons with Šidák adjustment (p < 0.05). Colors indicate domestication status as shown in (A).

### Photosynthesis and Quantum Yield

In week 0 (no difference between treatments; all plants were well-watered), maximum photosynthetic rate, estimated as the steady-state rate of CO_2_ assimilation at 800 PPFD (A_max_), was significantly higher in wild accessions relative to domesticated accessions (t_(31.3)_ = -3.256, p = 0.007) according to Tukey-adjusted tests for pairwise comparisons, indicating greater photosynthetic productivity in wild plants compared to domesticated plants under non-stressed conditions (Fig. 2A, Table S3). In weeks 1-2, when drought-treated plants did not receive any water, A_max_ was lower for plants in the drought treatment compared to the control treatment, and there was no significant effect of domestication status (wild, feral, or domesticated) on the reduction in A_max_ due to drought (Fig. 2A; Table S3). After rewatering (week 3), wild plants in the drought treatment had significantly greater recovery in A_max_ compared to domesticated plants in the drought treatment (t_(60.5_) = -3.13, p = 0.0076), indicating higher physiological resilience to multi-week drought stress in wild accessions (Fig. 2A; Table S3). Further, in week 3, wild accessions that underwent drought fully recovered maximum photosynthetic rates upon rewatering, with A_max_ values not significantly different from the same accessions in the control treatment (t_(123)_ = 0.50, p = 0.616). In comparison, A_max_ in drought-stressed domesticated (t_(117)_ = 2.22, p = 0.028) and feral (t_(115)_ = 3.31, p = 0.001) accessions remained depressed relative to the respective accessions in the control treatment (Fig. 2A).

**Figure 2.**
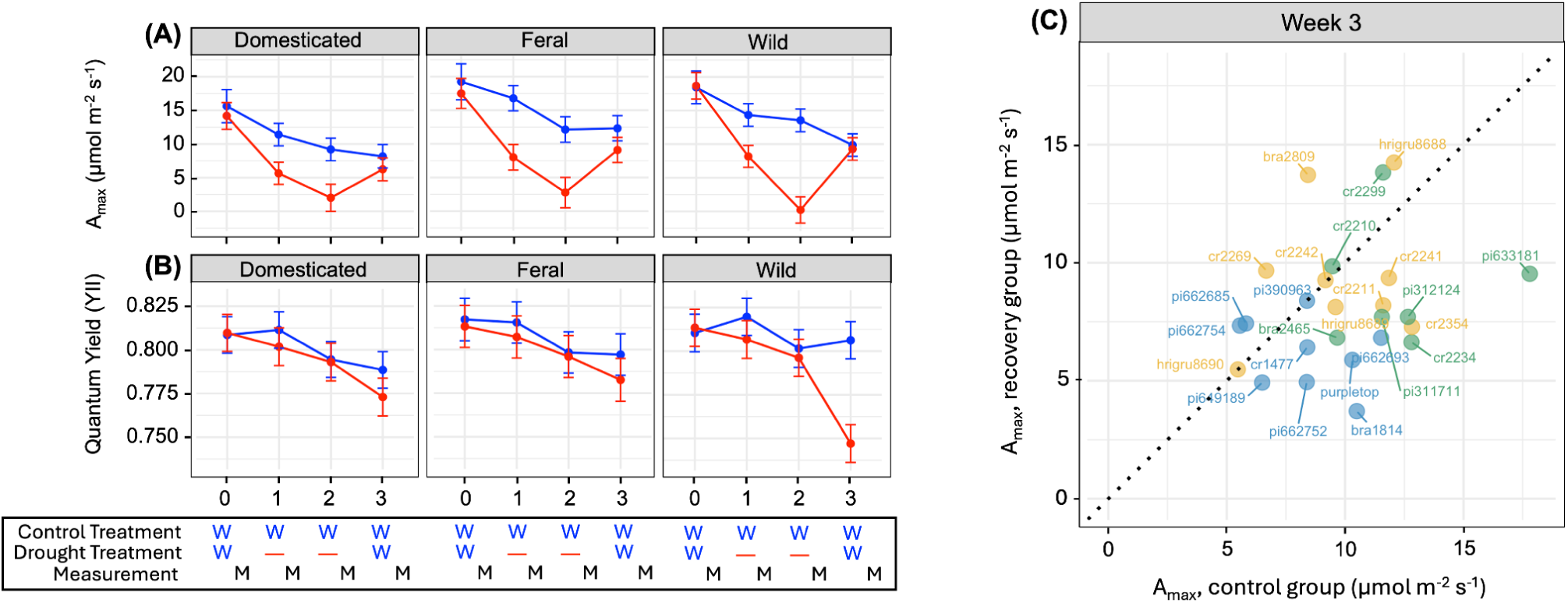
Leaf maximum photosynthesis (A) and quantum yield (B) over time. Week 0 = all plants were watered; week 1-2 = drought treatment received no water while control treatment received water; week 3 = all plants were watered. W indicates watering and M indicates measurements. (A) Steady-state CO_2_ assimilation rate at 800 PPFD in control (blue) vs. droughted (red) plants by domestication status. (B) Quantum yield of photosystem II (YII) in control vs. droughted plants; color scheme and experiment time series are the same as in (A). Points represent group means by domestication status; bars represent standard error (SE); solid lines connect group means. (C) Comparison of steady-state CO_2_ assimilation rate at 800 PPFD in the control group vs. the recovery group (droughted and re-watered) plants upon rewatering (week 3). The dashed line indicates a 1:1 linear relationship. Points indicate group means by accession (blue = domesticated, green = feral, gold = wild).

For plants in the drought treatment, the change in A_max_ from week 0-1 (shown as the initial slope in Fig. 2A) indicates the degree of sensitivity to drought, with a slope not significantly different from the plants in the watered treatment indicating a lack of drought sensitivity. The change in A_max_ from week zero to one was significantly lower in the drought than control plants for wild (t_(130)_ = 2.26, p = 0.025) and feral (t_(123)_ = 2.25, p = 0.026) plants, but not for domesticated plants (t_(132)_ = 1.65, p = 0.101), suggesting relatively lower sensitivity to drought in domesticated plants. However, it is important to note that in week 1, A_max_ was greater in wild and feral than in domesticated varieties, with A_max_ declining from week 0-1 in both watered and drought treatment domesticated plants (Fig. 2A). The change in A_max_ from week one to two was significantly different between the watered and drought treatment for wild plants (t_(128)_ = 5.74, p <0.0001), indicating sensitivity to drought. In contrast, the change in A_max_ from week one to two was not significantly different between the drought and control treatment for feral (t_(132)_ = 0.18, p = 0.855) and domesticated (t_(133)_ = 1.21, p = 0.229) plants.

The difference in A_max_ between weeks 2-3 (shown as the final slope in Fig. 2A) for drought treated plants indicates the degree of drought recovery (resilience). Drought treatment plants of all groups showed increased A_max_ between weeks 2-3 (positive slopes), indicating drought resilience. This trend differed by domestication group (Week × Treatment × Domestication interaction, χ² = 16.06, p < 0.001). Wild accessions fully recovered from drought, with no significant difference in A_max_ between drought and control plants in week 3 (t_(28.1)_ = 0.69, p = 0.497). In contrast, domesticated t_(30.4)_ = 2.12, p = 0.043) and feral (t_(38.2)_ = 3.07, p = 0.004) accessions still show reduced A_max_ after rewatering.

Wild plants in the drought treatment had the greatest increase in A_max_ between weeks 2-3, which was significantly steeper than plants in the control treatment (t_(115)_ = -7.27, p < 0.0001), indicating rapid recovery of assimilation rate upon rewatering. Thus, the wild varieties were both the most sensitive and the most resilient to drought.

Pre-dawn measurements of the quantum yield of photosystem II (YII) indicate overall leaf photosynthesis health. We found no significant difference in quantum yield between drought and control plants of all varieties in weeks 0, 1 and 2, (p-values all >0.05; Fig. 2B). In week 3 (following re-watering), wild plants showed significantly lower quantum yield for drought compared to control plants (t_(120)_ = 5.00, p < 0.0001, Fig. 2B), while there was no significant difference in quantum yield between drought and control plants for domesticated (t_(120)_ = 1.29, p = 0.199) and feral (t_(120)_ = 1.02, p = 0.309) varieties.

Notably, several accessions showed comparatively high A_max_ as well as full photosynthetic recovery upon rewatering in week 3, including HRIGRU 8688 (Wild; Caucasus) and CR 2299 (Feral; Colombia; Fig. 2C). These genotypes are likely both highly productive and drought resilient.

### Water Use Efficiency and Stomatal Conductance

In week 0, before the drought treatment, wild plants had significantly lower water use efficiency (WUE) than domesticated plants (t_(24.6)_ = 2.58, p = 0.042; Fig. 3A), but the difference from feral plants was not significant (t_(24.0)_ = 2.01, p = 0.131). Wild plants also had significantly greater stomatal conductance (g_sw_) than both domesticated (t_(24.6)_ = -5.36, p < 0.0001) and feral (t_(23.9)_ = -2.72, p = 0.031) varieties (Fig. 3B). In week 2 (two weeks post-drought), domesticated (t_(22.1)_ = 2.85, p = 0.0092) and wild plants (t_(20.1)_ = 3.66, p = 0.0016), had lower WUE in the drought compared to the control treatments, while there was no difference in WUE between drought and control for the feral plants (t_(31.6)_ = 0.20, p = 0.844). This indicates that none of the groups responded to the initiation of drought by becoming more water use efficient. Within the drought treatment in week 2, feral plants had significantly higher WUE than wild plants (t_(63.3)_ = 3.04, p = 0.0095). In week 3 (post re-watering), all varieties in the drought treatment showed an increase in WUE (domesticated: t_(5.71)_ = - 1.63, p = 0.16; feral: t_(6.23)_ = -3.38, p = 0.014; wild: t_(5.64)_ = -2.13, p = 0.08) and decrease in g_sw_ (domesticated: t_(7.34)_ = 1.56, p = 0.16; feral: t_(8.47)_ = 4.28, p = 0.002; wild: t_(7.01)_ = 1.68, p = 0.14; Fig. 3), with feral plants having significantly higher WUE and lower g_sw_ in the drought compared to the control treatment (Fig. 3). Within the control treatment, domesticated plants had significantly higher WUE than feral plants (t_(36.7)_ = 3.00, p = 0.013).

**Figure 3.**
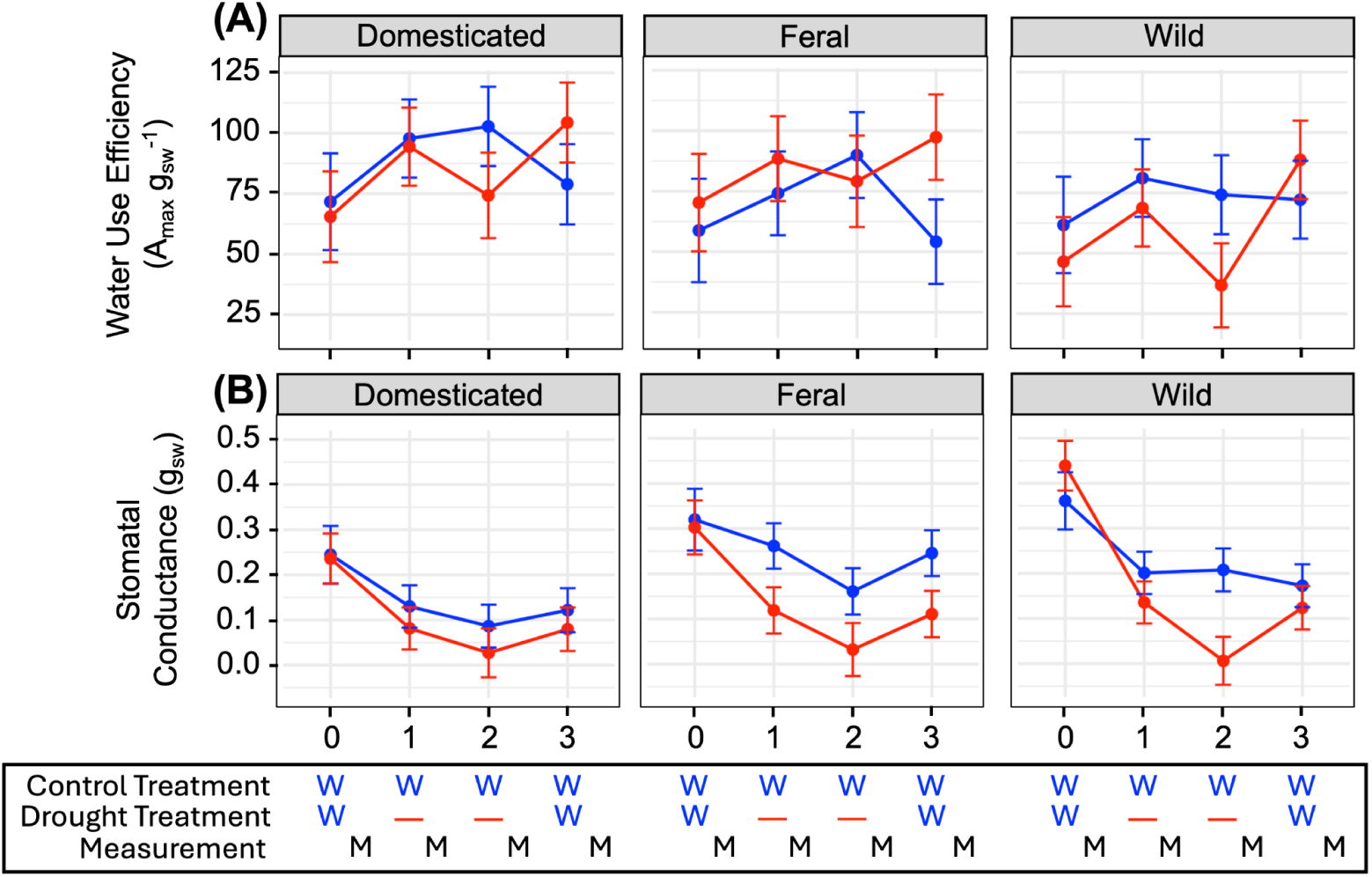
Water use efficiency and stomatal conductance timeseries. (A-B) Week 0 = all plants were watered; Week 1-2 = drought treatment received no water while control treatment received water; Week 3 = all plants were watered. W indicates watering and M indicates measurements. (A) Water use efficiency (WUE) in control (blue) vs. drought (red) treatments by domestication status. (B) Stomatal conductance to water (g_sw_) in control vs. drought treatments; color scheme and experiment time series are the same as in (A). Points represent group mean by domestication status; bars represent standard error (SE); solid lines connect group means.

Considering the change from week to week, WUE increased significantly overall (p < 0.0001) from week 0 to week 1, however, this increase occurred under both drought and control treatments (Fig. 3A), so this was not a drought-specific response. While there was a treatment × domestication effect when averaged across weeks (p = 0.0038), the rate of increase did not differ significantly for any domestication type across treatments (domesticated: t_(178)_ = -0.417, p = 0.678; wild: t_(187)_ = 0.40, p = 0.687; feral: t_(188)_ = -1.63, p = 0.106). Between weeks 1 and 2, domesticated plants showed a significant difference in slope between treatments (t_(238)_ = 2.46, p = 0.015), while wild (t_(236)_ = 1.55, p = 0.124) and feral (t_(237)_ = 1.05, p = 0.3) plants did not. However, the three-way interaction for this pattern was not significant (p = 0.6506). The treatment × domestication effect, averaged across the two weeks, was the most significant factor (p = 0.0009). Between weeks 2 and 3, drought affected WUE for all domestication types (week × treatment, p < 0.0001), and the effect was significant within each group tested separately (domesticated: t_(144)_ = -3.43, p = 0.0008; feral: t_(161)_ = -2.48, p = 0.0144; wild: t_(140)_ = -4.28, p < 0.0001), with wild plants showing the largest difference in slope between treatments, followed by domesticated and feral plants; however, there was no significant three-way interaction for week × treatment × domestication type (p = 0.483), indicating a trend rather than group differences. Domestication type (p = 0.0087), week × domestication type (p = 0.0015), and treatment × domestication type (p = 0.0075) were significant, indicating that the different domestication types differed in how WUE changed as the plants recovered from drought.

In week 0, wild plants had significantly higher baseline stomatal conductance (g_sw_) than domesticated (t_(24.6)_ = -5.36, p < 0.0001) and feral (t_(23.9)_ = -2.72, p = 0.031) plants. In week 1, there were significant differences between domesticated and feral plants in the control treatment (t_(58.6)_ = -4.19, p = 0.0003). In week 2, drought significantly reduced g_sw_ in feral (t_(8.02)_ = 3.52, p = 0.0078) and wild (t_(6.08)_ = 5.84, p = 0.0011) plants compared to the respective accessions in the control treatment. In week 3, feral plants in the drought treatment had significantly reduced g_sw_ compared to feral plants in the control treatment (t_(8.47)_ = 4.28, p = 0.0024), while domesticated and wild plants had statistically similar g_sw_ between treatments (domesticated: t_(7.34)_ = 1.56, p = 0.161 and wild: t_(7.01)_ = 1.68, p = 0.138).

Considering the rate of change in stomatal conductance, g_sw_ declined more rapidly in wild plants in the drought treatment compared to wild plants in the control treatment from week 0 to week 1 (t_(135)_ = 2.22, p = 0.028) and from week 1 to week 2 (t_(137)_ = 4.21, p < 0.0001). Between weeks 2 and 3, drought reduced g_sw_ for wild plants (t_(164)_ = -3.45, p = 0.0007), while domesticated and feral plants showed no significant differences in slope between drought and control treatments (domesticated: t_(168)_ = 0.118, p = 0.907 and feral: t_(184)_ = 0.785, p = 0.433).

### Taproot Biomass and Water Use Efficiency

Mean water use efficiency (WUE = A_max_ g ^-1^), calculated by accession and treatment over the 4-week time series, was positively associated with taproot dry biomass (R^2^ = 0.23, F(1,48) = 14.21, p = 0.00045; Fig. 4). Thus, plants with larger taproots were generally more water efficient.

**Figure 4.**
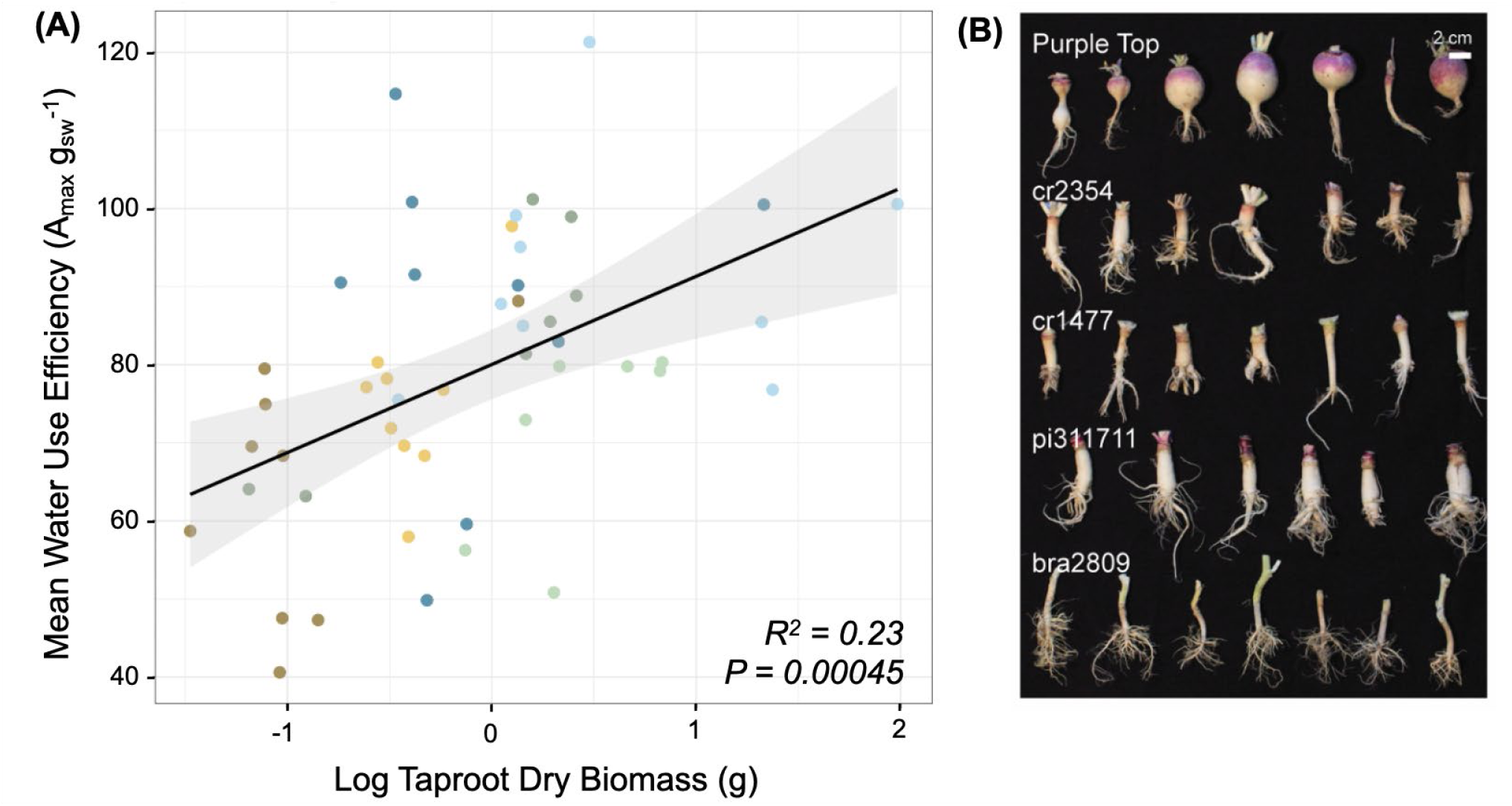
*B. rapa* taproot dry biomass association with water use efficiency. (A) Points represent group means by accession and treatment (control = light blue, light green, light gold; drought = dark blue, dark green, dark gold), colored by domestication status (domesticated = blue, feral = green, wild = gold). The solid line fits a linear regression and the gray shaded area indicates a 95% confidence interval band. Taproot dry biomass data were log transformed to normalize data distribution prior to regression modeling. (B) Representative variation in taproot biomass across domesticated (Purple Top, CR 1477), feral (PI 311711), and wild (CR 2354, BRA 2809) accessions.

## Discussion

Taking advantage of genetic variation in cultivated, feral, and wild *B. rapa* accessions, our results provide several insights into plant drought response across domestication status. We found there was substantial variation in drought resistance and resilience, with wild varieties often highly resilient in multiple dimensions of drought response (e.g., biomass retention, recovery of maximum CO_2_ assimilation rate) compared to crop and feral varieties, suggesting that conspecific wild relatives of crops can serve as valuable breeding material to face drought-related challenges. Specifically, we found that several accessions of *B. rapa* accumulated similar or even increased biomass under drought compared to well-watered conditions. Accessions with increased biomass in the drought treatment included the wild accessions CR 2354, CR 2242, and CR 2241 from the country Georgia. The fact that these accessions had greater biomass under drought than watered conditions, indicating overcompensation, is striking. Overcompensation to drought has been observed in multiple species, but our understanding of how this occurs is limited (Ke et al., 2025). In our study, we found the overcompensating accessions had slightly higher A_max_ and g_sw_ than the average accession; however, these differences were not statistically significant. Considering the adaptive landscape of these accessions, Georgia, and the Caucasus more broadly, is among the best documented regions for the presence of wild *B. rapa,* with populations there having split from domesticated forms an estimated 3,430–5,930 years before present (Sinskaja, 1969; McAlvay et al., 2021). This region has a highly diverse climate, with patterns of precipitation and drought that have varied widely over time (Elizbarashvili et al., 2025), which may facilitate the adaptation of wild *B. rapa* populations there to drought conditions.

There are several potential reasons that wild accessions had greater drought recovery than domesticated or feral forms. One important factor is that wild accessions in the drought treatment were shown to fully recover maximum CO_2_ assimilation rates (compared to the control treatment) after rewatering, with the combination of elevated A_max_ and recovery of A_max_ in wild accessions such as HRIGRU 8688 (Caucasus). This indicates higher resilience to multi-week drought stress in wild accessions than in feral and domesticated accessions, even with a significant decline in the quantum yield (YII) of wild plants after rewatering. Since YII is an indication of photosynthetic health and has been linked to xylem hydraulic failure (Brodribb et al., 2021), this suggests physiological tolerance for hydraulic failure as a mechanism for fast growth, potentially as part of a drought escape strategy. These results, together with reduced water use efficiency (WUE) in drought wild accessions, are consistent with drought escape through inefficient water use, rapid development, and early flowering, as had been previously reported in *B. rapa* (Franks, 2011).

While the drought escape strategy is usually viewed as completing the lifecycle before the onset of drought (Kooyers, 2015), our results suggest that this same combination of traits can also allow high drought resilience. Thus, drought occurring early in the growing season, or brief periods of drought, could both select for traits linked to rapid development that would allow plants to escape or quickly recover from drought. The idea that wild varieties use water prolifically to allow for rapid growth would also explain our finding that these varieties were both the most sensitive and the most resilient to drought.

A comparison of wild and crop varieties in our drought response experiment indicates that domestication diminished drought recovery in *B. rapa*, likely because crops were irrigated or grown in locations or during times with abundant precipitation. This finding is consistent with the idea that crops are often selected for increased yield in benign conditions of sufficient soil moisture and nutrients but lack stress tolerance compared to wild varieties. which are thus a good source of genes for increasing the ability of plants to respond to adverse conditions. It is important to note that because of trade-offs, increasing drought tolerance in crops could potentially decrease yield or other desirable crop traits (Renzi et al., 2022), so it is necessary to investigate the full effects of such stress response strategies. However, we identified several varieties that had both high drought resilience and high A_max_, indicating both high stress recovery and high productivity, likely due to rapid recovery of assimilation rate upon rewatering in combination with a fast growth strategy.

Increased drought resilience in wild relatives has been documented in other crop species such as cowpea (*Vigna unguiculata*) (De Meyer et al., 2026) and several cereal species (Toulotte et al., 2022). In one study, domesticated sunflower (*Helianthus annuus*) accessions experienced a 24% decline in biomass following drought, while the wild relatives did not experience statistically significant changes (Tran et al., 2024). Wild relatives of wheat vary widely in a range of drought-related traits, and different species and populations appear to use distinct strategies to conserve water, for example through reduced stomatal conductance or differences in root architecture (Merchuk-Ovnat et al., 2016; Pour-Aboughadareh et al., 2017). Crop wild relatives typically retain higher levels of genetic diversity than domesticated forms, and can preserve alleles that can confer resistance to drought, disease, and other challenges (Dempewolf et al., 2014; Brozynska et al., 2016; Zhang et al., 2017). This is often due to the domestication bottleneck effect, where diversity is constrained by generations of inbreeding, genetic drift, and selection, though the degree of these effects varies across crops (Von Wettberg et al., 2018; Tobón-Niedfeldt et al., 2022).

Because biomass allocation and growth strategies differ across crop types (leafy, oilseed, and root-hypocotyl), these types could differ substantially in drought response (Farooq et al., 2019). However, we did not find clear differences in drought response among the *Brassica* crop types assayed, possibly due to low replication for these groups (three leafy, two oilseed, and three turnip accessions). However, this does not preclude the possibility that drought response may depend on the harvested organ of the plant and may be apparent under different environmental conditions, so further studies on this topic would be useful.

Another key finding of our study is that plants with greater taproot dry biomass had greater instantaneous water use efficiency. This result is notable because it was previously thought that a large taproot could help plants tolerate or recover from drought by providing resources from this storage organ that plants could draw on when needed (Kalra et al., 2024), but this would not necessarily increase the efficiency of water use. In contrast, we found that taproot biomass was associated with greater WUE under drought conditions than plants with smaller taproots. This result could help to explain why crops with taproots were often able to quickly recover after drought, as was found in a recent meta-analysis (Dao et al., 2025), and shows the potential role of fleshy taproots in providing a hydraulic buffer for droughted plants and promoting water use efficiency.

Overall, the results of this study also contribute to an emerging theory of how domestication and ferality shape plant functional traits (Milla et al., 2015) and support the idea that feralization is not simply domestication in reverse (Niego and Benítez-Burraco, 2022). The contrast in drought response between the wild and feral accessions challenges assumptions that wild and feral accessions possess similar traits and can be treated together as uncultivated germplasm. Our findings suggest that these two groups diverge in drought response. These differences may be due to the different adaptive environments faced by wild and feral plants, with wild plants shaped by the need for tolerance to climate stress and feral plants adapting to disturbance and anthropogenic environments that may be more water and fertility-rich, such as farmland edges and suburban areas (Van Alfen, 2014; Mabry et al., 2023). In feral plants, these factors could lead to less conservative strategies around water use or less selective pressure on drought-resilient physiology (Mabry et al., 2023).

These findings suggest that feral populations and wild populations can contain suites of physiological traits that are very different from each other, affecting their utility for breeding. In addition, feral populations may also be adapted to other disturbance-heavy anthropogenic habitats such as roadsides, field margins, and waste areas. Further complicating generalizations of wild and feral populations is the fact that feral populations may arise with or without introgression with wild relatives, and feral *B. rapa* populations appear to have some proportion of ancestry from wild populations (McAlvay et al., 2021). This may partially explain some of the variation observed, such as the feral PI 312124 from Guatemala which had the 5^th^ highest biomass retention of the 25 accessions. Understanding these differences in physiological domestication and feralization syndromes may help guide pre-breeding efforts to identify resilience traits in non-domesticated populations (Lei et al., 2024).

Given the increasing intensity and duration of droughts in many areas due to climate change (IPCC, 2023), it is critical to understand how plants respond to and recover from drought (Xu et al., 2010; Fang and Xiong, 2015). Our findings demonstrate that evolutionary history and accession identity were important factors in determining drought response in *B. rapa*. The wild relatives CR 2354, CR 2242, and CR 2241 represent potential candidates for further investigation as a source of useful genetic variation due to their high biomass retention and rapid restoration of photosynthetic function following drought. Wild and feral accessions exhibited differentiation in terms of drought response, indicating that feralization does not necessarily imply a return to wild phenotypes. Wild and feral *B. rapa* relatives represent distinct genetic resources, and this evolutionary history should be considered in breeding programs to increase stress resistance and resilience.

## Supplementary data

Supplementary Figure S1. Experimental design.

Supplementary Figure S2. Soil moisture content (%) time series, by cultivar group.

Supplementary Figure S3. Biomass retention and crop type.

Supplementary Table S1. Trait table for 25 accessions of *Brassica rapa* (separate file).

Supplementary Table S2. Watering scheme for experimental control and drought treatments.

Supplementary Table S3. Estimated marginal means (± SE) for photosynthetic assimilation rate (A_max_) across weeks.

Supplementary Table S4. Estimated marginal means (± SE) for Quantum Yield (YII) across weeks.

Supplementary Table S5. Estimated marginal means (± SE) for Water Use Efficiency (WUE) across weeks.

Supplementary Table S6. Estimated marginal means (± SE) for Stomatal Conductance (g_sw_) across weeks

## Acknowledgments

We thank the horticultural staff of the Nolen Glasshouse (NYBG) for assistance with experiment setup and maintenance, and volunteer Khue Nguyen for assistance with cleaning root samples.

## Author Contributions

M.D., A.C.M., and C.T. conceptualized the study. A.C.M., A.M.B., K.I., A.E., and C.T. designed and executed the experiments, with data collection and analysis contributions from G.I.Q., G.A., R.W., L.R.V., M.L., R.M., I.G., R.S., J.B., L.C.D., E.M., G.M. and M.M. The manuscript was drafted by A.M.B., G.I.Q., S.J.F., and A.C.M. with significant contributions from G.M. and M.M. All authors contributed to manuscript review and revision.

## Conflict of Interest

The authors declare no conflict of interest.

## Funding

E.M., R.S., and J.B. were supported by the Science & Cultural Institutions Network (SCI Network) NYC. G.I.Q. was supported by the US National Science Foundation (Franks: IOS 2204374).

## Data availability

All data are available in the manuscript or deposited at https://osf.io/6z37u.

## Notes

https://osf.io/6z37u/overview

### Competing Interest Statement

The authors have declared no competing interest.

