## Supplementary Materials for "Drought Response and Recovery in Wild, Feral, and Domesticated *Brassica rapa*: Implications for Crop Climate Resilience"

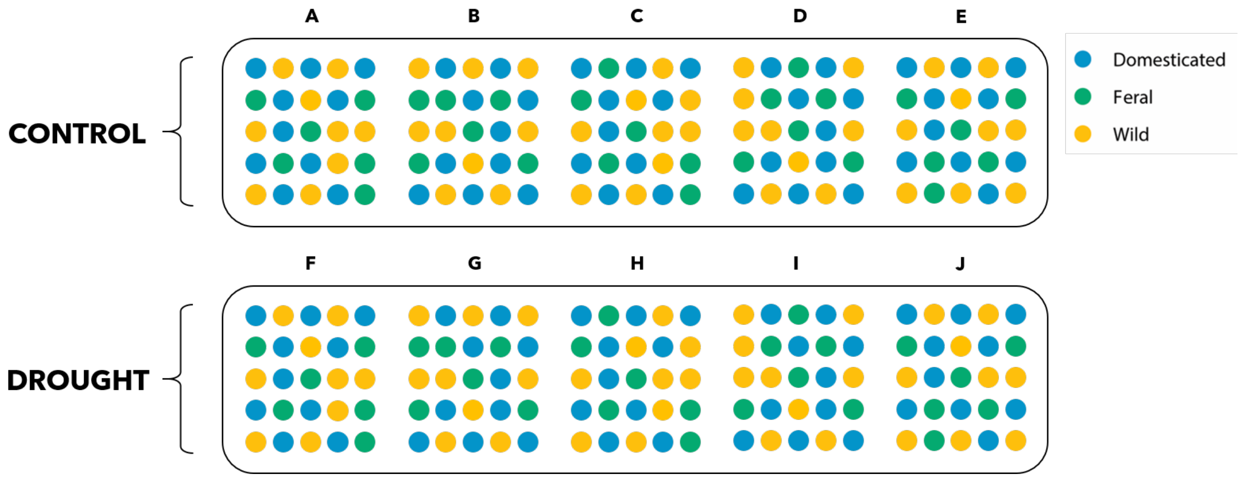


**Supplementary Figure S1.** Experimental design.

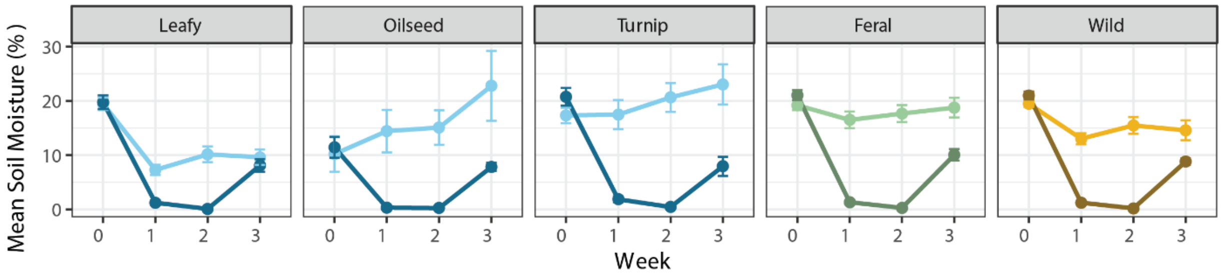


**Supplementary Figure S2.** Soil moisture content (%) timeseries, by cultivar group.

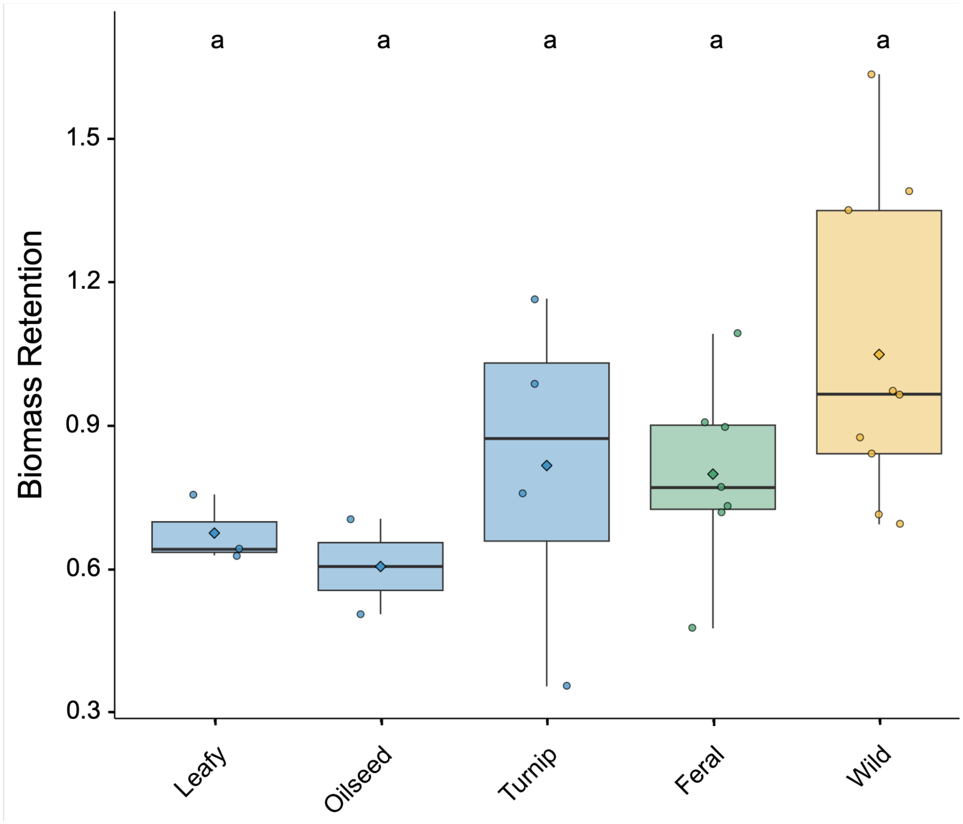


**Supplementary Figure S3.** Biomass retention and crop type. Mean and SE values are Leafy = 0.68 ± 0.16, Oilseed = 0.61 ± 0.19, Turnip = 0.82 ± 0.14, Feral = 0.80 ± 0.10, Wild = 1.1 ± 0.091.

**Supplementary Table S1.** Trait table for 25 accessions of *Brassica rapa* (separate file).

**Supplementary Table S2**. Watering scheme for experimental control and drought treatments.

| **Week** | **Control** | **Drought** |
|---|---|---|
| 0 | water | water |
| 1 | water | no water |
| 2 | water | no water |
| 3 | water | water |

**Supplementary Table S3.** Estimated marginal means (± SE) for photosynthetic assimilation rate ($A_{max}$) across weeks. Groups sharing a letter (within a week) are not significantly different (Sidak-adjusted, $\alpha = 0.05$).

| Week | Domestication Type | Treatment | Estimated Mean ($A_{max}$) | SE | 95% CI | Group |
|---|---|---|---|---|---|---|
| 0 | Domesticated | Control | 15.09 | 1.20 | [11.59 - 18.59] | ab |
| 0 | Domesticated | Drought | 14.76 | 1.03 | [11.70 - 17.83] | a |
| 0 | Feral | Control | 18.84 | 1.36 | [14.92 - 22.76] | ab |
| 0 | Feral | Drought | 17.68 | 1.17 | [14.29 - 21.08] | ab |
| 0 | Wild | Control | 18.89 | 1.22 | [15.22 - 22.56] | ab |
| 0 | Wild | Drought | 19.36 | 1.00 | [16.36 - 22.35] | b |
| 1 | Domesticated | Control | 11.35 | 1.38 | [6.85 - 15.85] | ab |
| 1 | Domesticated | Drought | 5.63 | 1.36 | [1.13 - 10.13] | a |
| 1 | Feral | Control | 16.67 | 1.47 | [12.09 - 21.26] | c |
| 1 | Feral | Drought | 7.95 | 1.49 | [3.34 - 12.55] | ab |
| 1 | Wild | Control | 14.11 | 1.38 | [9.61 - 18.61] | bc |
| 1 | Wild | Drought | 7.91 | 1.36 | [3.42 - 12.41] | ab |
| 2 | Domesticated | Control | 9.29 | 0.98 | [5.35 - 13.23] | bc |
| 2 | Domesticated | Drought | 2.44 | 1.23 | [-2.29 - 7.17] | ab |
| 2 | Feral | Control | 12.14 | 1.04 | [8.37 - 15.92] | cd |
| 2 | Feral | Drought | 3.44 | 1.29 | [-1.17 - 8.06] | ab |
| 2 | Wild | Control | 13.56 | 0.98 | [9.61 - 17.50] | d |
| 2 | Wild | Drought | 1.18 | 1.21 | [-3.60 - 5.96] | a |
| 3 | Domesticated | Control | 8.21 | 0.77 | [6.12 - 10.29] | ab |
| 3 | Domesticated | Drought | 6.14 | 0.74 | [4.11 - 8.16] | a |
| 3 | Feral | Control | 12.22 | 0.83 | [9.95 - 14.48] | c |
| 3 | Feral | Drought | 8.87 | 0.83 | [6.60 - 11.13] | ab |
| 3 | Wild | Control | 9.83 | 0.73 | [7.84 - 11.82] | bc |
| 3 | Wild | Drought | 9.37 | 0.72 | [7.42 - 11.33] | bc |

**Supplementary Table S4.** Estimated marginal means (± SE) for Quantum Yield (YII) across weeks. Groups sharing a letter (within a week) are not significantly different (Sidak-adjusted, $\alpha = 0.05$).

| Week | Domestication Type | Treatment | Estimated Mean (YII) | SE | 95% CI | Group |
|---|---|---|---|---|---|---|
| 0 | Domesticated | Control | 0.81 | 0.00 | [0.80 - 0.82] | a |
| 0 | Domesticated | Drought | 0.81 | 0.00 | [0.80 - 0.82] | a |
| 0 | Feral | Control | 0.82 | 0.00 | [0.81 - 0.83] | a |
| 0 | Feral | Drought | 0.81 | 0.00 | [0.80 - 0.82] | a |
| 0 | Wild | Control | 0.81 | 0.00 | [0.80 - 0.82] | a |
| 0 | Wild | Drought | 0.81 | 0.00 | [0.80 - 0.82] | a |
| 1 | Domesticated | Control | 0.81 | 0.00 | [0.80 - 0.83] | a |
| 1 | Domesticated | Drought | 0.80 | 0.00 | [0.79 - 0.81] | a |
| 1 | Feral | Control | 0.82 | 0.00 | [0.80 - 0.83] | a |
| 1 | Feral | Drought | 0.81 | 0.00 | [0.80 - 0.82] | a |
| 1 | Wild | Control | 0.82 | 0.00 | [0.81 - 0.83] | a |
| 1 | Wild | Drought | 0.81 | 0.00 | [0.80 - 0.82] | a |
| 2 | Domesticated | Control | 0.80 | 0.01 | [0.78 - 0.81] | a |
| 2 | Domesticated | Drought | 0.79 | 0.01 | [0.78 - 0.81] | a |
| 2 | Feral | Control | 0.80 | 0.01 | [0.78 - 0.82] | a |
| 2 | Feral | Drought | 0.80 | 0.01 | [0.78 - 0.81] | a |
| 2 | Wild | Control | 0.80 | 0.01 | [0.78 - 0.82] | a |
| 2 | Wild | Drought | 0.80 | 0.01 | [0.78 - 0.81] | a |
| 3 | Domesticated | Control | 0.79 | 0.01 | [0.76 - 0.81] | ab |
| 3 | Domesticated | Drought | 0.77 | 0.01 | [0.75 - 0.80] | ab |
| 3 | Feral | Control | 0.80 | 0.01 | [0.77 - 0.83] | b |
| 3 | Feral | Drought | 0.78 | 0.01 | [0.75 - 0.81] | ab |
| 3 | Wild | Control | 0.81 | 0.01 | [0.78 - 0.83] | b |
| 3 | Wild | Drought | 0.75 | 0.01 | [0.72 - 0.77] | a |

**Supplementary Table S5.** Estimated marginal means (± SE) for Water Use Efficiency (WUE) across weeks. Groups sharing a letter (within a week) are not significantly different (Sidak-adjusted, $\alpha = 0.05$).

| Week | Domestication Type | Treatment | Estimated Mean (WUE) | SE | 95% CI | Group |
|---|---|---|---|---|---|---|
| 0 | Domesticated | Control | 59.39 | 6.96 | [32.31 - 86.46] | ab |
| 0 | Domesticated | Drought | 61.85 | 5.58 | [33.57 - 90.14] | b |
| 0 | Feral | Control | 58.61 | 7.74 | [30.98 - 86.24] | ab |
| 0 | Feral | Drought | 58.07 | 6.05 | [32.89 - 83.25] | ab |
| 0 | Wild | Control | 51.49 | 7.36 | [21.86 - 81.13] | ab |
| 0 | Wild | Drought | 42.98 | 5.56 | [14.50 - 71.47] | a |
| 1 | Domesticated | Control | 93.59 | 10.12 | [62.81 - 124.37] | a |
| 1 | Domesticated | Drought | 97.93 | 10.05 | [67.23 - 128.63] | a |
| 1 | Feral | Control | 70.61 | 10.97 | [38.27 - 102.95] | a |
| 1 | Feral | Drought | 88.32 | 11.06 | [55.83 - 120.80] | a |
| 1 | Wild | Control | 82.43 | 10.08 | [51.75 - 113.10] | a |
| 1 | Wild | Drought | 65.21 | 9.96 | [34.71 - 95.72] | a |
| 2 | Domesticated | Control | 100.99 | 8.71 | [75.81 - 126.17] | b |
| 2 | Domesticated | Drought | 67.65 | 10.46 | [38.14 - 97.16] | ab |
| 2 | Feral | Control | 82.75 | 10.06 | [54.23 - 111.26] | b |
| 2 | Feral | Drought | 80.14 | 11.75 | [47.39 - 112.90] | ab |
| 2 | Wild | Control | 74.40 | 8.72 | [49.15 - 99.64] | b |
| 2 | Wild | Drought | 32.73 | 10.30 | [3.42 - 62.03] | a |
| 3 | Domesticated | Control | 88.46 | 9.50 | [57.08 - 119.85] | a |
| 3 | Domesticated | Drought | 107.57 | 9.42 | [76.18 - 138.95] | a |
| 3 | Feral | Control | 60.57 | 9.93 | [28.85 - 92.28] | a |
| 3 | Feral | Drought | 101.00 | 9.93 | [69.28 - 132.71] | a |
| 3 | Wild | Control | 73.62 | 9.29 | [42.25 - 104.98] | a |
| 3 | Wild | Drought | 98.48 | 9.18 | [67.17 - 129.80] | a |

**Supplementary Table S6.** Estimated marginal means (± SE) for Stomatal Conductance ($g_{sw}$) across weeks. Groups sharing a letter (within a week) are not significantly different (Sidak-adjusted, $\alpha = 0.05$).

| Week | Domestication Type | Treatment | Estimated Mean ($g_{sw}$) | SE | 95% CI | Group |
|---|---|---|---|---|---|---|
| 0 | Domesticated | Control | 0.28 | 0.06 | [-0.02 - 0.58] | ab |
| 0 | Domesticated | Drought | 0.27 | 0.05 | [-0.12 - 0.65] | a |
| 0 | Feral | Control | 0.38 | 0.06 | [0.10 - 0.66] | ab |
| 0 | Feral | Drought | 0.33 | 0.05 | [-0.01 - 0.66] | a |
| 0 | Wild | Control | 0.42 | 0.06 | [0.12 - 0.72] | ab |
| 0 | Wild | Drought | 0.47 | 0.05 | [0.08 - 0.86] | b |
| 1 | Domesticated | Control | 0.13 | 0.03 | [0.03 - 0.24] | a |
| 1 | Domesticated | Drought | 0.08 | 0.03 | [-0.03 - 0.19] | a |
| 1 | Feral | Control | 0.26 | 0.03 | [0.16 - 0.37] | b |
| 1 | Feral | Drought | 0.12 | 0.03 | [0.01 - 0.23] | ab |
| 1 | Wild | Control | 0.20 | 0.03 | [0.09 - 0.31] | ab |
| 1 | Wild | Drought | 0.13 | 0.03 | [0.02 - 0.24] | ab |
| 2 | Domesticated | Control | 0.10 | 0.02 | [0.02 - 0.18] | ab |
| 2 | Domesticated | Drought | 0.04 | 0.03 | [-0.06 - 0.13] | ab |
| 2 | Feral | Control | 0.17 | 0.02 | [0.09 - 0.25] | bc |
| 2 | Feral | Drought | 0.04 | 0.03 | [-0.05 - 0.14] | ab |
| 2 | Wild | Control | 0.22 | 0.02 | [0.14 - 0.30] | c |
| 2 | Wild | Drought | 0.02 | 0.03 | [-0.08 - 0.12] | a |
| 3 | Domesticated | Control | 0.11 | 0.02 | [0.03 - 0.19] | a |
| 3 | Domesticated | Drought | 0.06 | 0.02 | [-0.02 - 0.14] | a |
| 3 | Feral | Control | 0.23 | 0.02 | [0.15 - 0.31] | b |
| 3 | Feral | Drought | 0.09 | 0.02 | [0.02 - 0.17] | a |
| 3 | Wild | Control | 0.16 | 0.02 | [0.08 - 0.23] | a |
| 3 | Wild | Drought | 0.10 | 0.02 | [0.03 - 0.18] | ab |
